# The cell envelope of *Thermotogae* suggests a mechanism for outer membrane biogenesis

**DOI:** 10.1101/2022.09.22.508938

**Authors:** Danielle L. Sexton, Ameena Hashimi, Lloyd Sibanda, Polina Beskrovnaya, Tao Huan, Elitza I. Tocheva

## Abstract

The presence of a cell membrane is one of the major structural components defining life. Recent phylogenomic analyses have supported the hypothesis that the last bacterial common ancestor was likely a diderm. Yet, the mechanisms that guided outer membrane (OM) biogenesis remain unknown. Thermotogae is an early-branching phylum with a unique OM, the toga, previously shown to form 2-dimensional arrays of β-barrel trimers. Here we use cryo-electron tomography to characterize the *in situ* cell envelope architecture of *Thermotoga maritima*, proteomics and lipidomics to identify the protein and lipid composition of the toga, and bioinformatics to assess the distribution of the major toga components across the phylum. We show that the toga is composed of multiple Ompα and β-barrel homologs that represent a highly diverse bipartite OM-tethering system. We further reveal the presence of membrane microdomains (∼200nm) in the toga that are enriched in phosphatidylethanolamine (PE) lipids required to support the type 4 pilus and the BamA transmembrane complexes. Together, our results highlight a toga-like structure as a possible intermediate between monoderm and diderm cell envelope transitions.

## Introduction

The bacterial cell envelope is a diverse and complex structure that provides a barrier between the cell and its environment and facilitates interactions between the two (1). The presence of one (monoderm) or two membranes (diderm) surrounding the cell affords a way to classify bacteria by their cell envelope architecture. In monoderm bacteria, exemplified by *Bacillus subtilis*, the cell is encapsulated by a single lipid bilayer and a thick layer of peptidoglycan (PG). Diderm bacteria, such as *Escherichia coli*, have an inner and an outer membrane, separated by a thin layer of PG (1). Many bacteria, however, deviate from the canonical monoderm and diderm architectures. For example, Cyanobacteria are diderm with a thick layer of PG (2), while Mollicutes lack an OM and PG altogether(3). Furthermore, members of the diderm phylum Deinococcus-Thermus have an IM, thick layer of PG, an OM lacking lipopolysaccharides (LPS), and a proteinaceous surface layer (S-layer) (4, 5). The bacterial cell envelope can further vary in lipid and protein composition, and the presence of a polysaccharide capsule. Studies into the diversity of the bacteria cell envelope structure and function can give valuable insight into its adaptive roles in various environment, as well as provide deeper understanding of the evolutionary paths of its major components.

The inner (IM, or cytoplasmic CM) and outer membranes (OM) in bacteria are structurally and functionally different. The IM contains a-helical proteins and sustains a proton gradient, while the OM is rich in β-barrel proteins and facilitates the free diffusion of small molecules into and out of the periplasm. In particular, the Omp85/BamA superfamily of OM proteins (OMPs) have distinct β-barrel tertiary structures and are essential for the proper incorporation of other β-barrel proteins in the OM. Broadly conserved membrane lipid components include glycolipids known as lipopolysaccharides (LPS), found in the outer leaflet of typical OMs. Another major feature in diderm bacteria is the presence of a molecular tethering system. In the Terrabacteria (early branching phyla), this is formed by the OmpM/SlpA superfamily. These proteins contain an N-terminal S-layer homology (SLH) domain that binds PG, a long coiled-coil domain spanning the periplasm, and a C-terminal β-barrel component embedded in the OM (6, 7). The domain architecture of these proteins is conserved across most diderm Terrabacteria, and evolutionary scenarios involving SlpA/OmpM losses have been proposed as a mechanism for diderm to monoderm transition in Firmicutes (7). Overall, the presence of β-barrel proteins, particularly those of the BamA family, an OM-tethering system, and LPS glycolipids are considered hallmarks of archetypical diderm bacteria and are often used to predict the presence of an OM. Recent advances in metagenomics have expanded our appreciation of microbial diversity and provide an ample opportunity to study how and when major structural features of the bacterial envelope have evolved.

Thermotogae is a member of the clade Terrabacteria, which includes other early-branching phyla such as Cyanobacteria, Firmicutes, Actinobacteria, Deinococcus-Thermus, Chloroflexi and the Candidate Phyla Radiation (CPR) (8, 9). Frequently placed near the base of the bacterial tree of life, Thermotogae are mostly anaerobic, hyperthermophilic bacteria that stain Gram-negative. Although their cell envelope consists of an IM and a thin layer of PG, due to their unique OM, they are considered atypical diderms (10). The OM, referred to as ‘toga’, is formed by an extended proteinaceous sheath that can become distanced from the IM at the cell poles and as such is credited for their signature appearance. Genomic analyses have revealed the absence of key genes for LPS biosynthesis (*lpx)* and LPS transport (*lpt*) (11-13) indicating that they lack LPS. Nevertheless, members of the phylum encode for a BamA homolog that has been used as a proxy for the presence of an OM (11). Another distinct feature of Thermotogae is the production of several unusual membrane lipids, including membrane-spanning lipids, widely present in archaea but rarely found in bacteria. These membrane-spanning C_30_ and C_32_ diabolic acids contain tetraether, tetraester, or mixed ether-ester linkages with phosphoglycerol head groups on one end and glycerol molecules on the other (14). Membrane-spanning lipids have been shown to organize into lipid monolayers with increased rigidity and decreased permeability compared to phospholipid bilayers (15), a possible adaptation to hyperthermal environments.

Insights into the structure of the cell envelope of Thermotogae mainly come from studies on *Thermotoga maritima*, the best characterized member of the phylum. It was proposed that Ompα and Ompβ (previously OmpA/B) were the major structural components of the toga (16-19). Ompα was shown to contain a 45-nm long coiled-coil domain (17) and a conserved SLH domain that bound to PG-associated secondary cell wall polymers (19, 20). A gene (*ompβ*) identified in an operon with *ompα* was predicted to have high β-sheet content and was proposed to encode for the β-barrel forming the trimers (19). More recently, Ompα and Ompβ were suggested to act analogously to the OmpM/SlpA tethering system that, in other early-branching phyla, is encoded by a single protein (5, 7). Additional major structural components of the toga include the well-characterized hydrolases xylanase, amidase, and xylosidase, used by the bacteria to degrade complex carbohydrates in the environment and generate metabolic energy (21).

Here we use a multidisciplinary approach including cryo-electron tomography (cryo-ET), MS-based proteomics and LC-MS-based lipidomics to characterize the structure and composition of the cell envelope of *T. maritima*. Our tomograms reveal the *in situ* organization of the cell envelope in 3-dimensions to macromolecular resolution (∼3 nm), compositional analyses define the protein and lipid components of the IM and toga, and homology searches show the distribution of major toga and toga-associated proteins throughout the Thermotogae phylum. We discuss the evolutionary implications of our findings among other early-branching phyla and propose a mechanism for OM biogenesis that implicates a toga-like structural intermediate.

## Results

### Cell envelope architecture of T. maritima

Cryo-electron tomography (cryo-ET) was used to visualize intact *T. maritima* cells (Fig. 1, n=35). Tomograms revealed that the cell envelope consisted of an IM, a thin layer of PG and a toga (Fig. 1A). Along the lateral sides of the cell, the periplasmic space was consistently ∼70±10 nm wide while it varied at the poles (∼450±100 nm). The IM appeared as a 4-nm thick monolayer, whereas the toga featured two distinct arrangements: 1) a 4-nm thick monolayer (Fig. 1B, purple), and 2) a 7-nm thick bilayer (Fig. 1D, light blue). Density profiles of the cell envelope at the monolayer and bilayer sections further revealed single and double toga peaks, respectively (Fig. 1C, E). Top (xy) views of the toga at the tips of cells displayed an extended sheath with a 2-dimensional pattern (Fig. 1F, Supplementary Data Fig. S1, Movie S1). 2D Fourier transforms and subtomogram averaging further confirmed the presence of porin-type proteins arranged in a trimer and forming a hexagonal array with 9.8 nm lattice spacing (Fig. 1F). Previous studies using freeze-etching calculated the lattice spacing to be 11 nm (16), a difference likely be due to differences in the methods used. The bilayers patches of the toga were 7-nm thick, ∼100-200 nm in length, and void of surface patterns. Based on these observations, we concluded that the bilayers were membrane patches.

**Figure 1.**
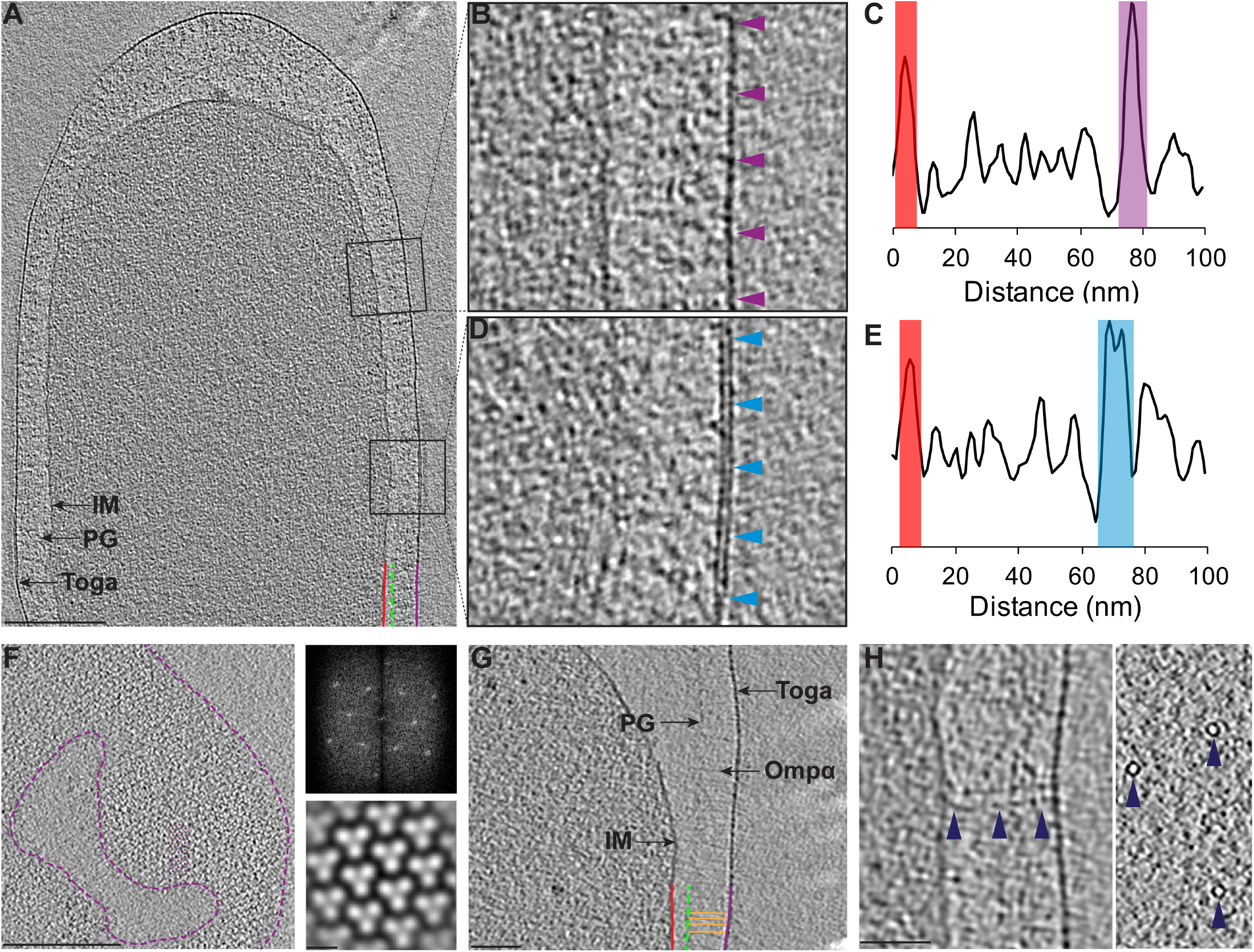
Ultrastructure of the *T. maritima* cell envelope. **A)** A 20-nm thick slice through a cryotomogram. The cell envelope is composed of an inner membrane (IM, red), a thin layer of peptidoglycan (PG, green), and the toga (purple). The toga is consistently spaced (∼70 nm) from the IM along the lateral sides of the cell and balloons out at the poles. **B)** Enlarged view of the toga monolayer (purple arrows) and **C)** corresponding density profile. **D)** Enlarged view of the toga bilayer (light blue arrows), and **E)** corresponding density profile. **F)** Top view of the toga shows extended areas of repeating trimers bound by purple dashed lines. A small patch highlighting the array pattern is highlighted with purple dots (left). 2D Fourier transform (top right) and Subtomogram average (bottom right) revealed a hexagonal array with with 9.8 nm lattice spacing. **G)** Numerous copies of Ompα (orange) connecting the PG to the toga. **H)** Side (left) and top (right) views of T4P (dark blue arrows) connecting the IM to the toga. Scale bar 200 nm (A, F), 50 nm (G, H).

### Additional ultrastructural features

A continuous layer of Ompα densities (Fig. 1G) was observed spanning ∼50 nm underneath the toga and connecting the toga to the PG (Fig. 1E) consistent with previous studies (17, 19). Numerous tubular structures ∼80-nm long and 10 nm in diameter, consistent with a type IVa pilus (T4P) trans-envelope complex (22, 23) were observed spanning the periplasm and connecting the IM to the toga (Fig. 1H, Fig. S1, Movie S1). Additional densities, likely xylanases, were widely distributed along the exterior surface of the toga (Fig. S2A). Dividing cells revealed simultaneous invagination of the IM, PG-Ompα layer, and the toga, with putative FtsA and FtsZ filaments polymerized at ∼16 nm underneath the IM at division sites (Fig. S2B).

### Lipid composition and labeling of the toga

To confirm the presence of lipids in the toga, we performed lipidomic analyses and lipid labeling experiments. Lipidomics of the IM and toga revealed that both fractions broadly contained the same lipid species, though enrichment of certain lipids was detected in each fraction (Table 1). Ether and ester linked lipids were equally distributed between the IM and the toga and so were membrane spanning lipids, mono and di-glycosylated diacylglycerol (DAG), and acyl ether glycerol (AEG) species, including those with atypical deconyl-diGlu head groups (Supplementary Data Table S1). Phosphatidylethanolamine (PE) lipids, short free fatty acids (FAs) (C_10_, C_12_, C_13_), and select digalactosyldiacylglycerols (DGDGs) were enriched in the toga fraction whereas glycerol-linked diabolic acids (glycerol-DAs) were enriched in the IM. One species of DGDG (with C_16_C_19_) was present exclusively in the toga fraction (Table 1). All detected lipids above the set threshold and their relative abundancies are listed in the Supplementary Data Table S1.

Labeling with lipid dyes further revealed the presence of distinct puncta in the toga. Since PE lipids were enriched in the toga, we used a fluorescently-labeled PE derivative OG-DHPE to label cells. Fluorescent puncta, ∼200 nm in diameter, were observed in the toga of exponentially and stationary growing cells, consistent with our tomograms (Fig. 2A-C). Weak uniform staining of the IM was also observed, consistent with the presence of PE lipids in the IM. To improve the resolution of the fluorescence signal and identify additional detail of the puncta, we used super-resolution microscopy (Fig. 2D-F). In exponentially growing cells, the puncta were ∼120 nm in diameter and variably distributed in the toga, again consistent with the size and distribution of the membrane patches observed in tomograms. Images of the toga in stationary phase also revealed that the PE lipids formed puncta (Fig. 2G-I). Super resolution microscopy further revealed circular patterns with diameters of ∼700 nm and <200 nm. (Fig. 2J-L). The distribution of the OG-DHPE in the toga substantiates with the presence of lipids forming distinct microdomains.

**Figure 2.**
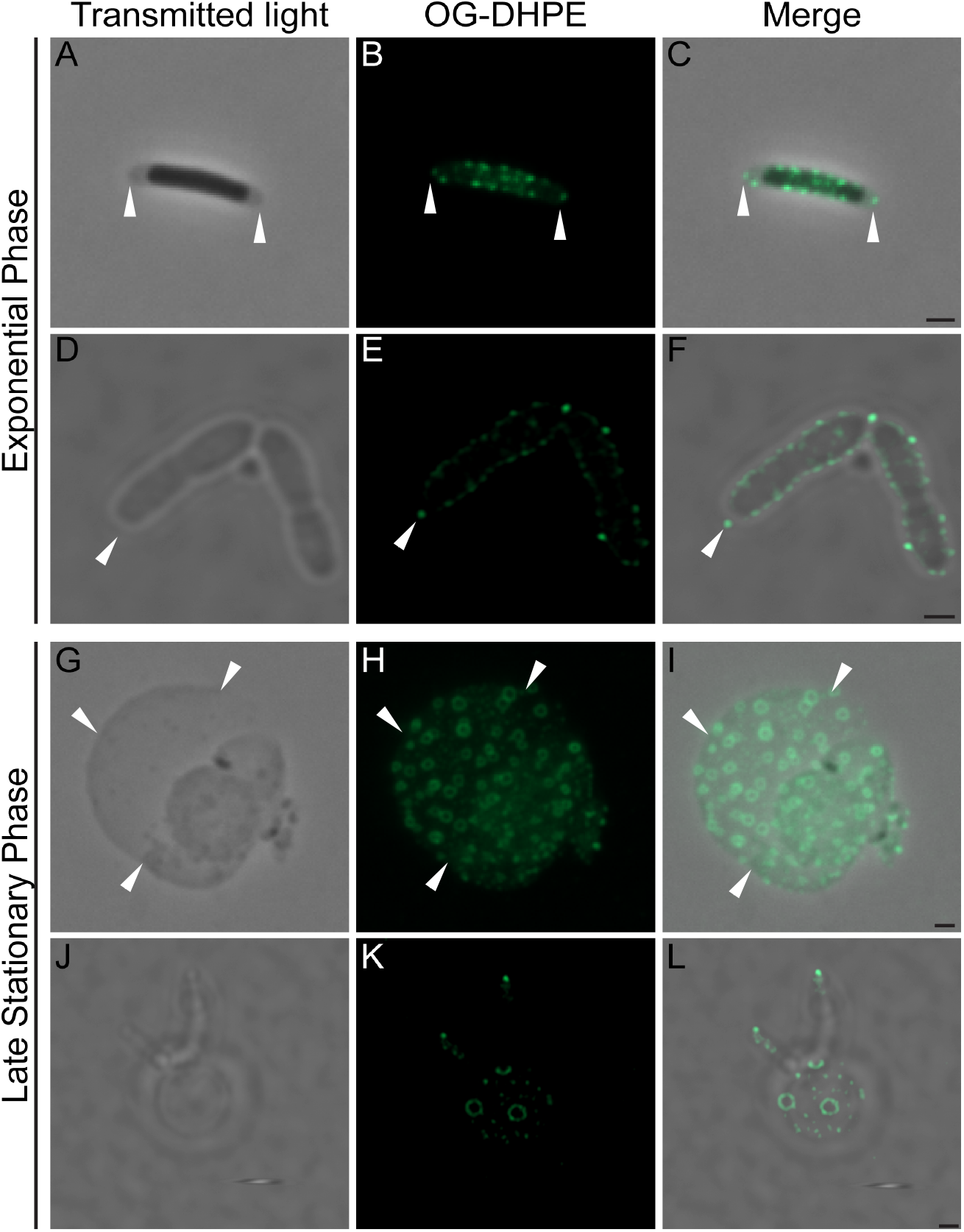
Detection of lipid domains in the toga of *T. maritima*. The fluorescent dye Oregon Green-DHPE (OG-DHPE) was used to stain cells in exponential (**A-F**) and stationary (g-l) growth phases. Cells were imaged with phase contrast light microscopy (**A-C, G-I**) to clearly detect areas of the toga that were separated from the IM (white arrows). Super resolution confocal microscopy (**D-F, J-L**) was used to improve the resolution from 300 to 100 nm. Images revealed distinct puncta within the toga of exponentially growing cells. Cells in stationary phase revealed circular patterns of PE lipids in the toga (∼700nm), as well as small puncta (<100nm) below the resolution limit of our experimental setup. Scale bar, 1 µm.

### Protein composition and structure prediction of the toga

MS-based proteomics was used to characterize the protein content of the IM and the toga. Table 2 lists the 15 most-abundant proteins in the toga, which notably included three Ompα homologs (Ompα1-3) with similar sizes (∼45kDa). Tertiary structure prediction revealed that all three homologs had a conserved SLH domain and formed extended coiled-coils (Table 2). Using AlphaFold2 Multimer, the three Ompα homologs were predicted to form trimers ∼50 nm in length, similar to the distance between the toga and the PG.

BamA and Ompβ were readily detected in the toga and both were predicted to form β-barrels (Table 2). Furthermore, three additional, highly-abundant proteins: R4NP97, G4GDI2, R4NSJ1 (∼45-50 kDa) were predicted to have Sec secretion signals and form β-barrels (Table 2). All five predicted β-barrel proteins had extensive hydrophobic surfaces along the central portion of their surfaces and charged belts at the top and bottom (Fig. S3). AlphaFold2 Multimer predicted that Ompβ, R4NP97 and G4GDI2 could form trimers with pore sizes ∼45-50 Å, consistent with our subtomogram averages (Fig. 1F). The models had high per-residue confidence (predicted local-distance difference test [pLDDT]) values, an indicator for high accuracy of the model (Fig. S1). The distribution of hydrophobic and charged amino acids on the exterior faces of the predicted structures could facilitate packing of these putative β-barrel proteins into the hexameric lattice observed in the toga. Compared to SlpA (trimer from *D. radiodurans*), Ompβ was predicted to have extremely negatively-charged external surfaces, which could be important for interactions with surface-associated carbohydrate hydrolases or determine the surface properties of the cells.

The most abundant protein detected in the toga fraction was xylanase A, an enzyme that hydrolyses xylan into xylose (Table 2). Other enzymes, likely surface-exposed and involved in polysaccharide hydrolysis, such as maltodextrin glucosidase and β-xylosidase (24), were also enriched in the toga, along with the OM pore forming protein PilQ and pilins, components of the T4P. Numerous T4P were observed in tomograms (Fig. 1H), consistent with the MS-based proteomics. Top abundant proteins also included an S-layer-like array protein (R4P1L1) and an uncharacterized protein with a helical structure (R4P086). Supplementary Data Table S2 lists all detected proteins in the toga with 95 % confidence.

### Phylogenetic distribution of major toga proteins

Homologs of abundant toga-associated proteins were highly-conserved among multiple members of the family Thermotogaceae, however, only BamA, Ompα1, and Ompα2 were found conserved in all members of the phylum (Fig. 3). Using a two-step approach, conserved SLH domains were identified in 152 proteins across the Thermotogae, 144 of which also had a coiled-coil domain (Supplementary Data Table S3). This approach confirmed that all Ompα homologs shared overall domain architecture, despite significant (up to 70 %) sequence variation in the coil-coiled region. Notably, Ompβ was poorly conserved, with homologs detected only in the genera Thermotoga and Thermosipho. Furthermore, Ompα2 and Ompβ were found in an operon solely in the genus Thermotoga, and in an operon with either Ompα1 or Ompα2 in the genus Thermosipho, contrary to previous proposals (16, 25). Of the uncharacterized proteins with predicted β-barrel tertiary structure, G4FDI2 was the most well-conserved (with some exceptions in Pseudothermotoga family) and homologues of the predicted β-barrel proteins R4NP97 and R4NSJ1 were missing from the Kosmotogaceae family, with the latter only consistently found in the genus Thermotoga. Together with the trimer prediction, these results suggest that BamA, Ompα1, Ompα2, G4FDI2 and R4NP97 are the major conserved components of the toga among members of the phylum.

**Figure 3.**
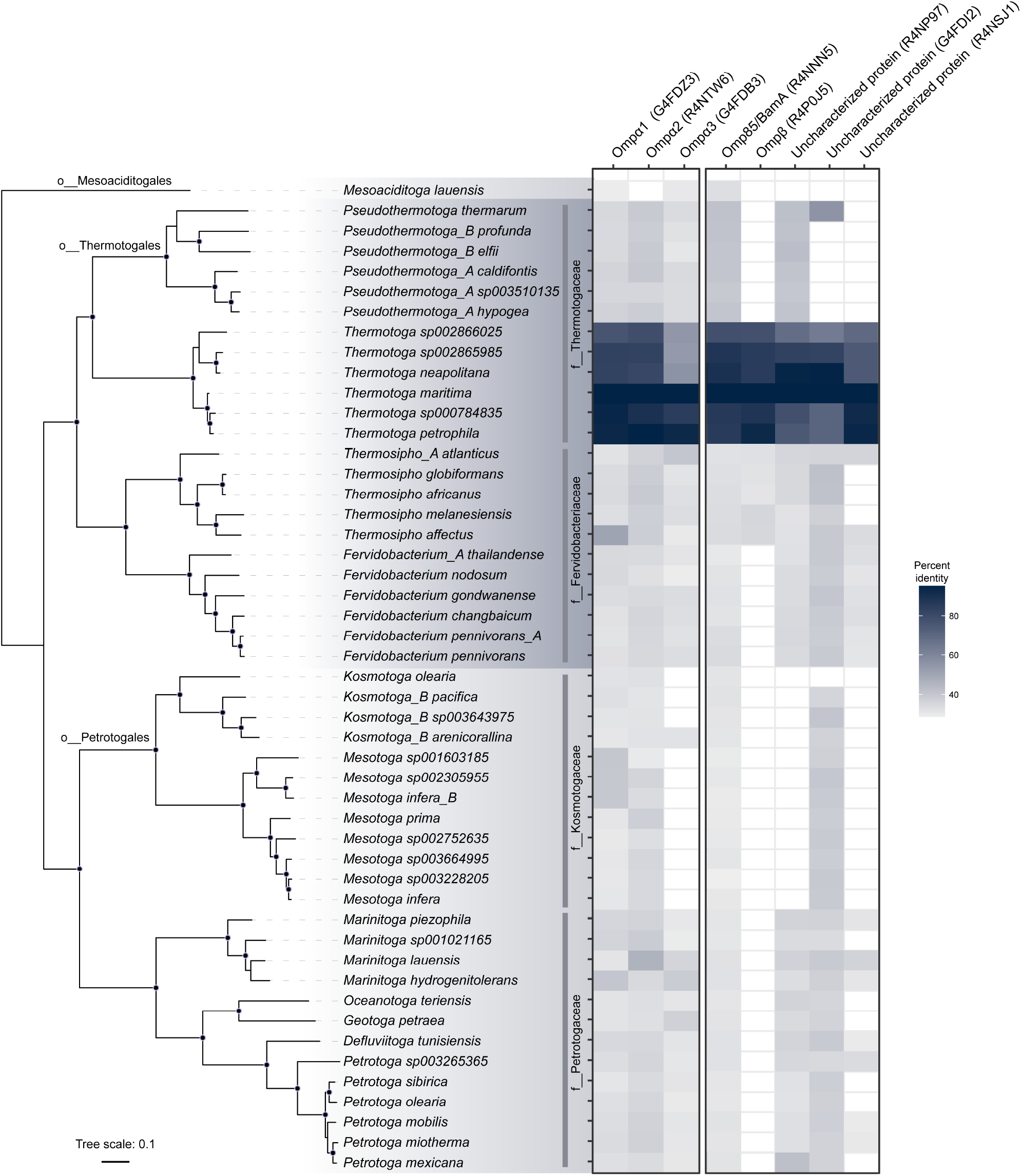
Conservation of toga-associated proteins across the Thermotogae phylum. Hits are shaded by % amino acid similarity to *T. maritima* proteins detected with mass spectrometry. Phylogenetic tree was constructed using a concatenated alignment of 120 single-copy marker genes and IQ-TREE with 1000 ultrafast bootstraps, and is shaded by taxonomic order. Coprothermobacter was used as an outgroup. Species (italic) and family (f___) names are listed as shown.

## Discussion

In this study, we characterize the unique cell envelope and toga composition of *T. maritima*. Previous studies showed that the toga was composed of a 2D-repeating array of porin trimers (16), and suggested Ompβ as the major porin forming the toga (19, 25). Our tomograms confirm that the toga is composed of an extended proteinaceous sheath of trimer porins (Fig. 1F), however, in addition to Ompβ, MS-based proteomics detect three other proteins of similar sizes, structure and abundance in the toga (Table 2). The expression of four proteins with predicted β-barrel structures: Ompβ, R4NP97, G4FDI2, and R4NSJ1, indicates that these proteins act as major structural components of the toga. All four are predicted to have similar surface potentials, mostly hydrophobic in the middle and charged on the periphery, similar to the monomeric BamA and trimeric SlpA (Fig. S3). Since the surface potentials of the β-barrels could not be used to predict whether the proteins assembled, multimeric structural prediction was used to provide further insight. Interestingly, only Ompβ, R4NP97 and G4FDI2 were predicted to form trimers. Since the latter two are best conserved in other species, we hypothesize that they are the major structural components of extended proteinaceous sheath. Deletion mutants of these proteins in *T. maritima* and assessment of the toga integrity would ultimately determine their roles.

Previous bioinformatic analyses showed that the OM-tethering system OmpM/SlpA is conserved in all known diderm Terrabacteria phyla and is encoded by a single protein, with the Thermotogae being a notable exception (7). Here the tethering system is encoded by two separate proteins: Ompα (containing the SLH domain and a coiled-coil domain) and a β-barrel protein. We show that all three Ompα homologs are found in the *T. maritima* toga and homology searches indicate that they are highly conserved among all Thermotogae species. While we detect Ompβ in the toga, our analyses show that it is poorly conserved beyond the genera Thermotoga and Thermosipho. In addition, Ompβ is encoded in an operon with Ompα2 only in the genus Thermotoga, making its role less significant than previously suggested. The three other proteins we identified with predicted β-barrel structures were not found in an operon with any of the Ompα homologs, though all three are distributed throughout the phylum. The need for different porins may have evolved as an adaptation to extreme environments since different expression patterns could provide the toga with modified properties. With the exception of BamA, none of the β-barrel proteins in Thermotogae have homologs in other phyla, which further suggests a unique evolutionary pathway of their tethering system. Since the existence of a toga has been documented in all Thermotogae classes (26-35) and Thermotogae are known for their highly dynamic genomes (36), we conclude that members have developed a tethering system with high degree of plasticity. Notably, structural conservation of the β-barrel fold is highly preserved among bacteria as it is more related to function than protein sequence.

In addition to the 2D-repeating array of porin trimers, our tomograms reveal the presence of lipid bilayers in the toga. The IM and toga contain the same lipid species as previously shown for *D. radiodurans* (37) and large proteins (LptF/LptG) with homology to ABC permeases have been suggested to represent a highly divergent system that relays lipids to the toga (13). Together with the presence of unique membrane-spanning lipids, Thermotogae represent a distinctive model for membrane formation. The use of fluorescently-labeled lipids identified FMMs with enriched PE lipids in the toga. The presence of FMMs in bacteria has been well-documented and these domains were suggested to have similar roles to the lipid rafts found in eukaryotic cells (38). FMMs concentrate proteins that function in related cellular processes such as sensor kinases involved in signal transduction, secretion systems facilitating membrane trafficking and other regulatory complexes (39). For example, cholesterol-rich microdomains in *Helicobacter pylori* were shown to contain high concentrations of T4SS cagPA1 complexes (40). Similarly, the observed FMMs in *T. maritima* could provide support for the membrane-embedded T4P and BamA complexes. Other properties of enrichment of certain lipids such as cholesterol and PE have been linked to a decrease in membrane fluidity, which could stabilize macromolecular complexes and sustain membrane function in thermal environments (41, 42).

### Evolutionary implications

Major questions about the biogenesis and evolution of the cell envelope, particularly the OM of diderm bacteria, remain unresolved. Due to its unique cell envelope and ancient origin, the Thermotogae phylum provides valuable insight into OM biogenesis. In particular, our study reveals that the toga of *T. maritima* is composed of extended 2D protein arrays and small membrane patches. As such, we propose that the toga represents a model for a structural intermediate between monoderm and diderm cell envelopes. Current phylogenomic evidence suggests that the last bacterial common ancestor (LBCA) was a diderm(9, 11). Yet, only three mechanistic models have been put forward describing how a diderm cell envelope could have evolved *de novo*: 1) Cavalier-Smith proposed that two half-cells fused of to make a protocell with a double membrane envelope (43), 2) Blobel theorized that protein-protein interactions on the surface of lipid vesicles could have mediated the formation of a diderm ancestor (44), and 3) Tocheva proposed that an ancient sporulation-like event could have transitioned a monoderm primordial cell into a diderm LBCA (45).

Based on our studies of *T. maritima*, here we propose another model for OM biogenesis where a toga-like structure could have served as a structural intermediate (Fig. 4A). Briefly, due to the seemingly simpler architecture, we hypothesize that the primordial cell was monoderm (Fig. 4A, step 1). Over time, the lipid membrane was strengthened by additional surface layers composed of protein and possibly PG (step 2). Blebbing parts of the primordial membrane accumulated and eventually integrated into the surface protein layer, forming a structure analogous to the toga (step 3). Notably, a key property of the surface layer would have been the ability to stabilize lipid molecules. Over time, continuous build-up of the lipid molecules in the surface layer would have given rise to a second membrane (step 4). We may never know the nature of the primordial surface layer; however, likely candidates are the immunoglobulin (Ig)-like domain-containing proteins. Ig-like domain-containing proteins have been shown to assemble into S-layers in monoderm and diderm bacteria, as well as in archaea (5, 46, 47). In *T. maritima*, an S-layer-like array protein with five Ig-like domains was detected as a major protein in the toga, suggesting it can also interact with lipids and β-barrel proteins (Table 1, Fig. 4C). These observations highlight the diverse properties and ancient origin of the Ig-like domains.

**Figure 4.**
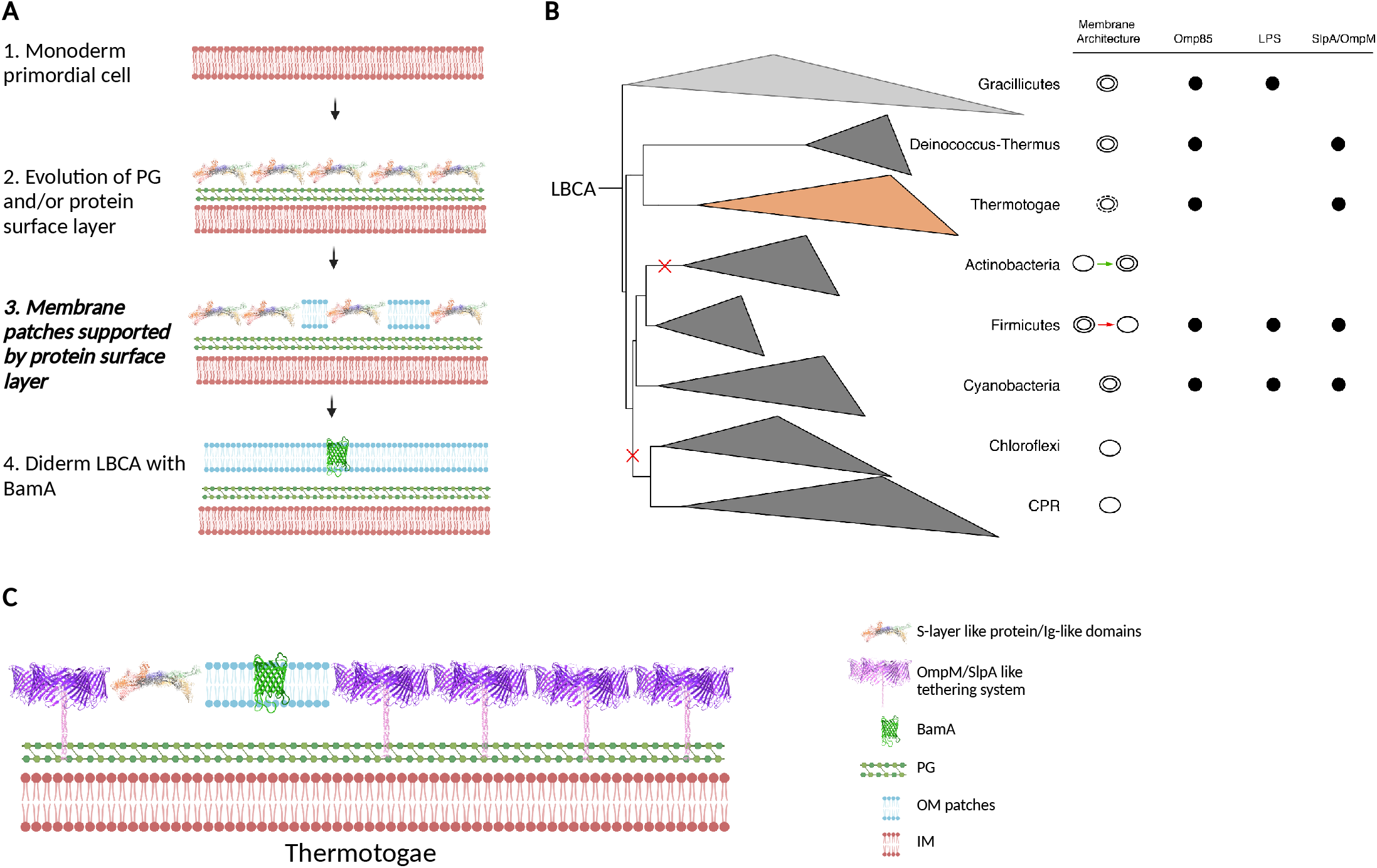
The toga as a model for OM biogenesis. **A)** Major evolutionary steps outlining a proposed mechanism for OM biogenesis: 1) the primordial cell was likely monoderm, 2) PG and/or a protein surface layer evolved to support the bilayer of the primordial cell, 3) the abundance of surface proteins was able to support small membrane patches (analogous to the toga, bold), and 4) over time, the membrane patches extended to form a second membrane. Since the β-barrel fold is ancient, BamA evolved concurrently with the OM. **B)** A phylogenetic tree of Terrabacteria depicting the major early-branching phyla (adapted from Witwinowski *et al*. 2022). Diversity in the early branching phyla could be explained with losses and gains of the OM as shown in red and green, respectively. Membrane architecture annotations are based on cryo-ET studies (except CPR, uncultured). Presence of OM signature features such as Omp85/BamA superfamily, LPS synthetic pathway, SlpA/OmpM tethering system, is indicated with black circles. **C)** Cell envelope diagram for Thermotogae shows major structural components pertinent to the OM biogenesis model (step 3, panel A).

Since BamA requires membrane support for proper function, our model suggests that BamA co-evolved with the diderm cell envelope, a notion with significant implications for OM biogenesis in Terrabacteria (Fig. 4B). Starting with a diderm LBCA, gains and losses of the OM could explain the diversity in cell envelope architectures. As noted previously, CPR and Chloroflexi likely lost their second membrane (48), while Actinobacteria lost it and eventually evolved a novel OM (49). It is now well-established that monoderm Firmicutes went through a diderm intermediate (50, 51) and lost their OM through a possible perturbation of their OM-tethering system (48). Deinococcus-Thermus and Cyanobacteria retained their OM and Thermotogae likely reverted to a toga intermediate.

## Materials and Methods

### Strains and growth conditions

*Thermotoga maritima* MSB8 was obtained from the DSMZ German Collection of Microorganisms and Cell Cultures and grown in DSMZ medium 1232 or TMB medium with 0.5% xylose (52). Liquid media were sterilized by passing through a 0.22 μ filter, then flushed with N_2_ gas. Cultures were manipulated in a BactronEZ anaerobic growth chamber (Sheldon Manufacturing Inc) with an atmosphere of 5 % H_2_, 10 % CO_2_, and 85 % N_2_ and incubated at 80 °C for 16 hrs to reach mid-exponential growth phase.

### Cryo electron tomography, data reconstruction, and subtomogram averaging

Bacterial cultures were mixed with 20-nm colloidal gold, loaded onto glow discharged carbon grids (R2/2, Quantifoil) and plunge-frozen using a Vitrobot Mark IV (ThermoFisher Scientific), maintained at room temperature and 70 % humidity, into liquid ethane-propane cooled to liquid nitrogen temperatures. Tilt series were collected using SerialEM (53) on a Titan Krios 300 kV transmission electron microscope (ThermoFisher Scientific) equipped with a Falcon III direct electron detector. Data collection conditions were -6 µm defocus, 120 e^-^/Å total dose, +/- 60° oscillations, with 1° tilt increments. Tilt series were aligned using IMOD, CTF corrected using CTFFind4 and novaCTF, and 3D reconstructions were calculated using IMOD with the back-weighted projection method (54-57). Density profiles were calculated using ImageJ (58). For subtomogram averaging of the toga, 1584 subvolumes were extracted and averaged with Dynamo (59). Alignment was refined 8 times without applying symmetry.

### Membrane labeling and fluorescence light microscopy

In order to detect lipids in the toga, we used the Oregon Green 488 1,2-Dihexadecanoyl-*sn*-Glycero-3-Phosphoethanolamine (OG-DHPE, ThermoFisher Scientific). OG-DHPE is a synthetic PE lipid analog coupled with a fluorophore that embeds into membranes with PE lipids. Since lipidomic analyses detected enrichment of the PE lipids in the toga, we expected that the toga will be brightly labeled with OG-DHPE compared to the IM. Prior to labeling, cells were grown for 16 hrs at 80 °C, washed three times with PBS pH 7.5 under anoxic conditions, and resuspended in PBS and incubated with 10 µg/mL OG-DHPE for 1 hr at room temperature. Cells were washed three times with PBS pH 7.5 and imaged with phase contrast and super-resolution fluorescence light microscopy using Zeiss Axio Examiner and Zeiss LSM 900 confocal microscope equipped with an Airyscan 2 detector. Images were collected and processed using Zen Blue 2.6 and 2.4 software respectively.

### Inner membrane (IM) and toga isolation

All membranes were isolated from a 2-g wet weight cell pellet from 1.5 L of culture following established protocols (60). Briefly, cells were resuspended in 10 mL lysis buffer (25 mM Tris pH 7.4, 5 mM ETDA, 500 mM NaClO_4_, 1 mM PMSF, 40 µg/mL lysozyme, 20 U/mL DNase) and passed two times through a homogenizer at 50 psi. Unlysed cells and large debris were removed by centrifugation at 4,000 *g* for 10 min. The supernatant, containing the whole membrane fraction, was isolated by ultracentrifugation at 100,000 *g* for 2 hrs. The membrane pellet was washed three times with 10 mL wash buffer (25 mM Tris pH 7.4, 5 mM ETDA, 500 mM NaClO_4_), and pelleted each time at 100,000 g for 1 hr. To separate the IM from the toga, the whole membrane fraction was resuspended in 200 µL wash buffer and layered on a sucrose gradient containing 1.2 mL of 70 % (w/w), 2 mL of 50 % (w/w), and 1.6 mL of 20 % (w/w) sucrose in wash buffer, and spun at 100,000 *g* for 2 hrs. Bands were removed from the top and were washed once with 20 mL wash buffer to remove excess sucrose.

### Lipid extraction and analysis

Whole membrane, IM, and toga fractions were analyzed for their lipid composition using liquid chromatography-mass spectrometry (LC-MS). Lipids were extracted in methanol:acetonitrile: water (2:2:1, vol:vol:vol) and methyl-tert-butyl ether (MTBE). Solvent was evaporated from the lipid containing fraction using vacuum concentration. The resulting dried residue was reconstituted in isopropanol:acetonitrile (1:1, vol:vol) and lipid profiling of cellular membranes was carried out using a reverse phase Acquity UPLC BEH C_18_ column (1.0 × 100 mm, 1.7 µm, Waters) on an Agilent 1290 Infinity II ultra-high performance liquid chromatography (UHPLC) system (Agilent Technologies) coupled with Bruker Impact II electrospray-ionization quadrupole time-of-flight (ESI-QTOF) mass spectrometer (Bruker Daltonics). Lipid identification was completed on MS-DIAL 4.7 tool based on mass accuracy, isotope ratio, retention time along with MS/MS similarity against publicly available libraries. MS-DIAL 4.7 can be downloaded at the PRIMe website (http://prime.psc.riken.jp/) (61). Raw lipidomics data were collected in positive and negative ion mode, after intensity correction and normalization (post quality control calibration). Quality control (QC) calibration was done to obtain more accurate fold changes: the relative standard deviation (rsd) of the QC injections (at the beginning, middle and end of the injection sequence) were below the threshold of 0.25. To measure enrichment of lipid species, fold changes (FC) of 0.67 and 1.5 were used as thresholds for the IM and toga, respectively. For example, FC were calculated as IM/toga ratio, thus FC of 0.67 indicated lipid enrichment in the toga, whereas FC of 1.5 indicated lipid enrichment in the IM fraction. Additional details of the lipidomics analysis can be found in the Supplementary Data.

### Protein identification and structure prediction

Fractions from the sucrose gradient containing the IM and toga were visualized on an SDS-PAGE gel and excised for subsequent digestion with trypsin for MS analysis. Tryptic peptides were analyzed on an Orbitrap Fusion Lumos Tribrid mass spectrometer (ThermoFisher Scientific) operated with Xcalibur (version 4.0.21.10) and coupled to a Scientific Easy-nLC 1200 system. Monoisotopic precursors were searched against the *T. maritima* MSB8 proteome (Uniprot ID 243274) using RawConverter (v1.1.0.18; The Scripps Research Institute). Peptides identified with a score having a confidence higher than 95 % were kept for further analysis. Additional details of the proteomics analysis can be found in the Supplementary Data. Protein localization was predicted using pSORTb (62), β-barrel prediction was done using BOMP and PRED-TMBB2 (63, 64), and tertiary structures were predicted using RoseTTAFold (65). Trimer structure predictions were performed using AlphaFold2 Multimer v1.0 (66) and the surface hydrophobicity and electrostatic charge for the highest ranked model were computed in ChimeraX (67). Per residue confidence plots (pLDDT) for all structures are available in Fig. S4.

### Phylogeny and sequence-based homology searches

To generate a phylogenetic tree of the phylum Thermotogae, the maximum-likelihood algorithm was used to extract an alignment of 120 conserved bacterial single-copy marker genes (68) from 63 available Thermotogae members using the Genome Taxonomy Database GTDB-Tk v1.4.0 software (69). The tree was constructed using IQ-TREE v2.1.4 with 1,000 ultrafast bootstraps and the substitution model LG+F+R6, as determined by ModelFinder (70, 71). Representative Coprothermobacterota genomes served as an outgroup. Publicly available phylogenetic tree data, created from the alignment of concatenated RNA polymerase subunits β, β′ and elongation factor IF-2 proteins rooted between Gracillicutes and Terrabacteria, was modified to represent early branching phyla (7). Both trees were visualized using iTOL (v4) (72).

To assess the genomes of Thermotogae members for the presence of homologs of toga-associated proteins detected with MS-based proteomics in *T. maritima*, we performed gene prediction using Prodigal on publicly available Thermotogae genomes from the GTDB to generate proteomes with locus tag information (40, 68). PSI-BLAST searches were performed on these proteomes using 4 iterations and an E value of 1e^-10^. To detect Ompα proteins, we used a two-step approach. First, we searched all Thermotogae genomes for proteins with an SLH domain using the Pfam profile PF00395. Next, we inspected all sequences with N-terminal SLH domains for the presence of a central coiled-coil segment using PCOILS (73). Since none of the identified proteins had a C-terminal β-barrel component, the presence of a β-barrel in an operon with the identified Ompα homologs was assessed manually.

## Acknowledgements

This project was supported by a Natural Sciences and Engineering Research Council of Canada Discovery Grant to EIT (RGPIN 04345). DLS was supported by a Natural Sciences and Engineering Research Council of Canada Postdoctoral Fellowship (546024) and AH was supported by a Natural Sciences and Engineering Research Council of Canada Post Graduate Scholarship (552674). We thank Dr. Florian Rossman and the High Resolution Macromolecular Cryo-Electron Microscopy facility at the University of British Columbia for assistance with microscope operation and tilt series acquisition. We also thank Dr. Laurent Brechenmacher at the Southern Alberta Mass Spectrometry (SAMS) Center.

## Supplementary Figures

**Figure S1.**
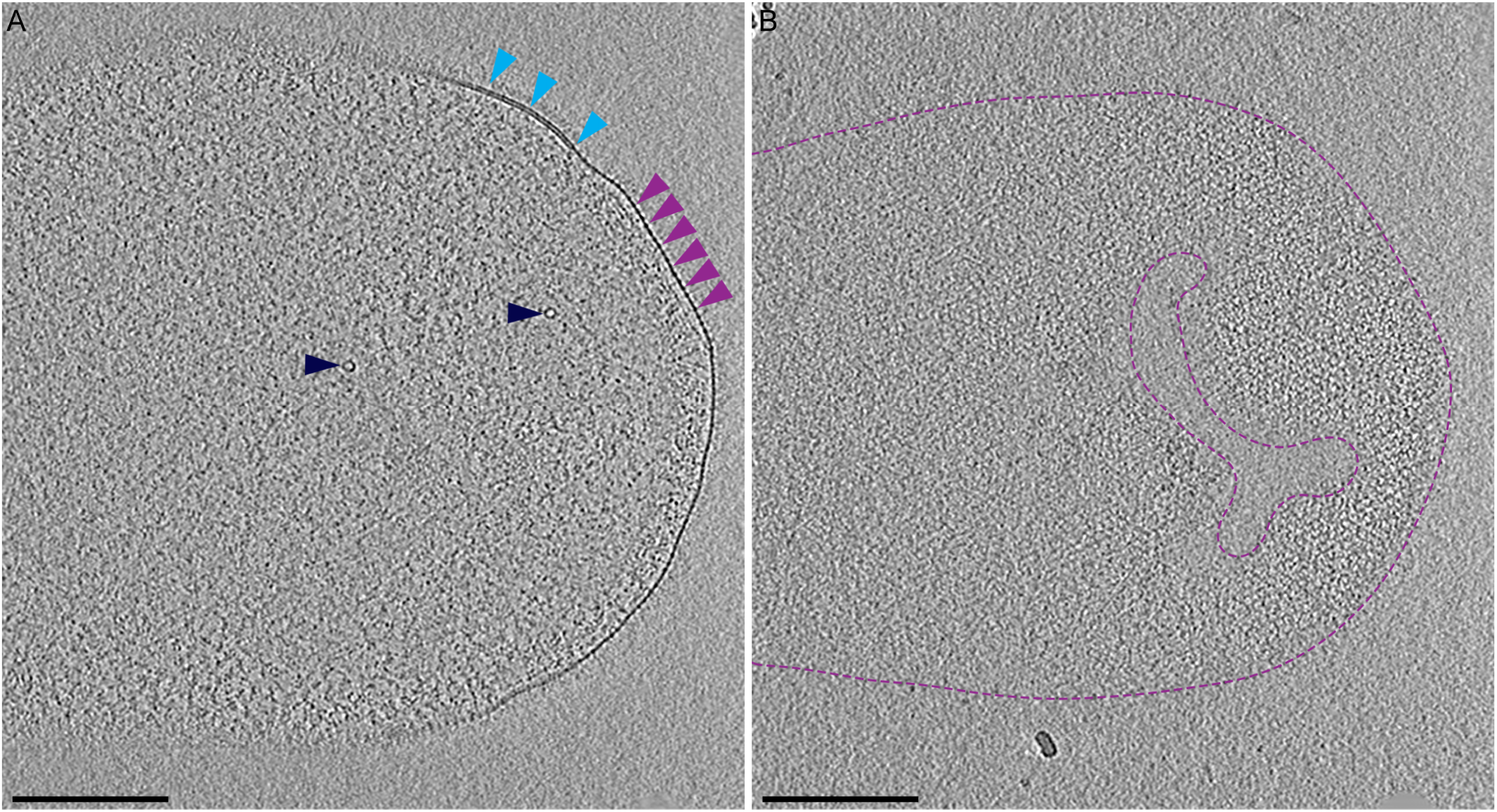
Highlights of *Thermotoga maritima* ultrastructure from Movie S1. **A)** A 20-nm thick section through the center of a cryotomogram shows a functional membrane microdomain (light blue arrows) and the repeating pattern of the protein monolayer in the toga (purple arrows). Top views of T4P (dark blue arrows) were observed embedded in the toga. **B)** A 20-nm thick tomographic slice revealed top view of the toga with the extended lattice of trimers (purple dots). The surface area covered in trimers is bound by purple dashed lines. Scale bar, 200 nm.

**Figure S2.**
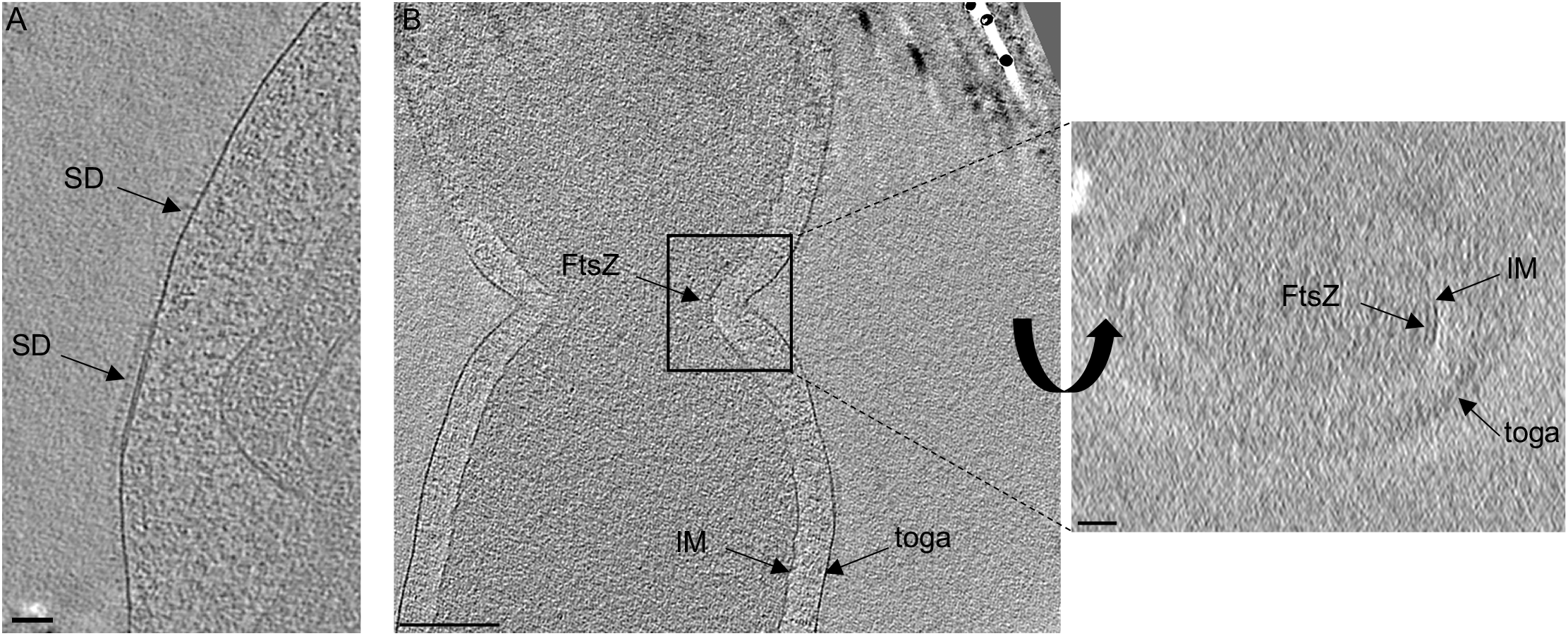
Additional ultrastructure in *Thermotoga maritima*. **A)** Surface densities (SD, likely composed of xylanase A, maltodextrin glucosidase and beta-xylosidase) were observed all along the cell surface and were more prominent on the membrane patches of the toga. **B)** A division site revealed that both the inner membrane (IM) and toga invaginate together during cell division. FtsZ was observed as a filament underneath the IM, ∼16 nm from the membrane (right panel).

**Figure S3.**
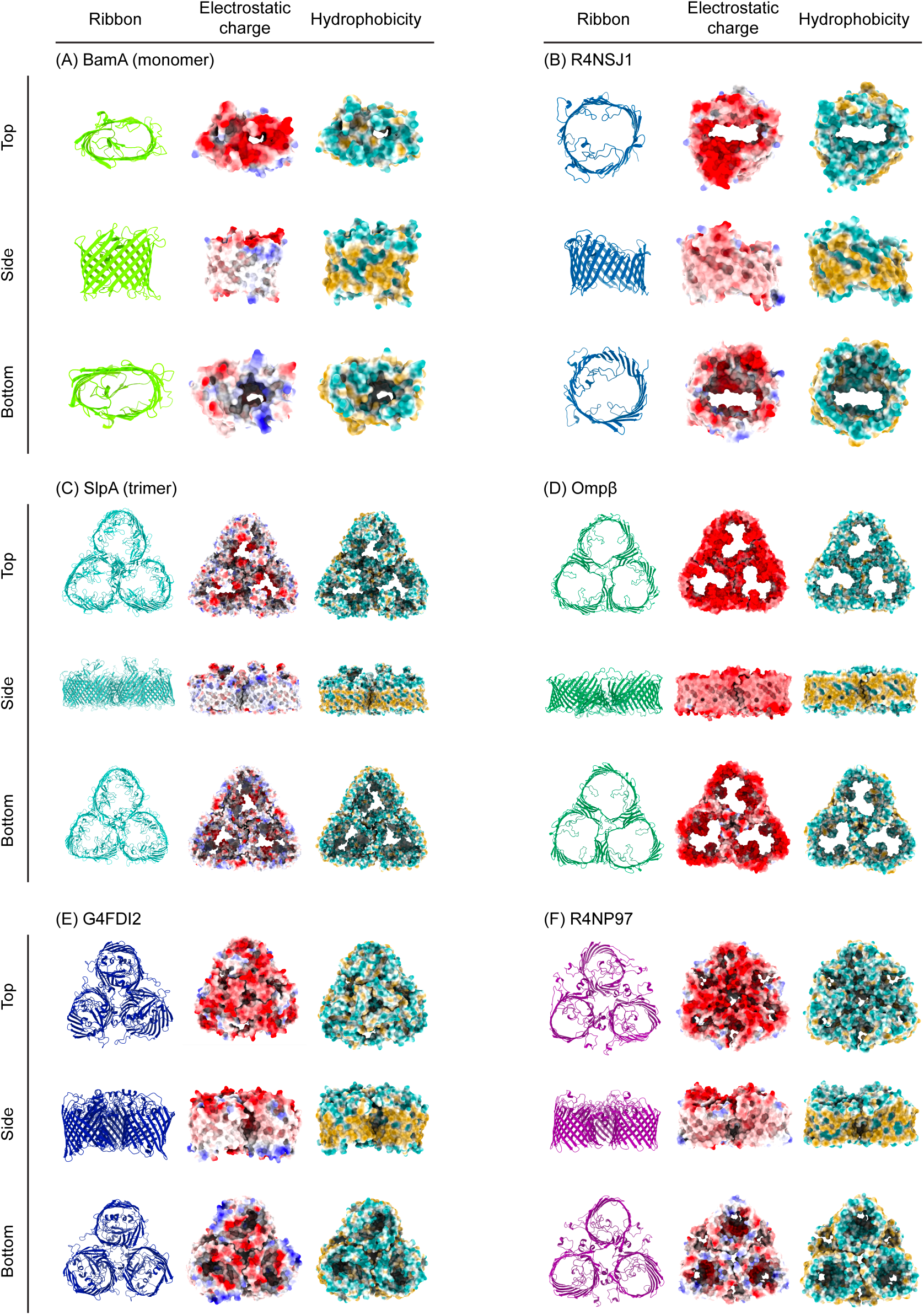
Predicted structures of putative β-barrel proteins in the toga. Trimeric protein structures for the β-barrel portion of all proteins were predicted using AlphaFold2 Multimer. For β-barrel proteins that did not form a trimer when modeled, the predicted structure of a single protein is depicted. This included **A)** the β-barrel portion of BamA, a known monomeric protein, and **B)** R4NSJ1. Predicted trimers included **C)** SlpA, a known trimeric protein from *Deinococcus radiodurans*, **D)** Ompβ, **E)** G4FDI2, and **F)** R4NP97. The electrostatic charge and surface hydrophobicity for each protein were modeled using ChimeraX.

**Figure S4.**
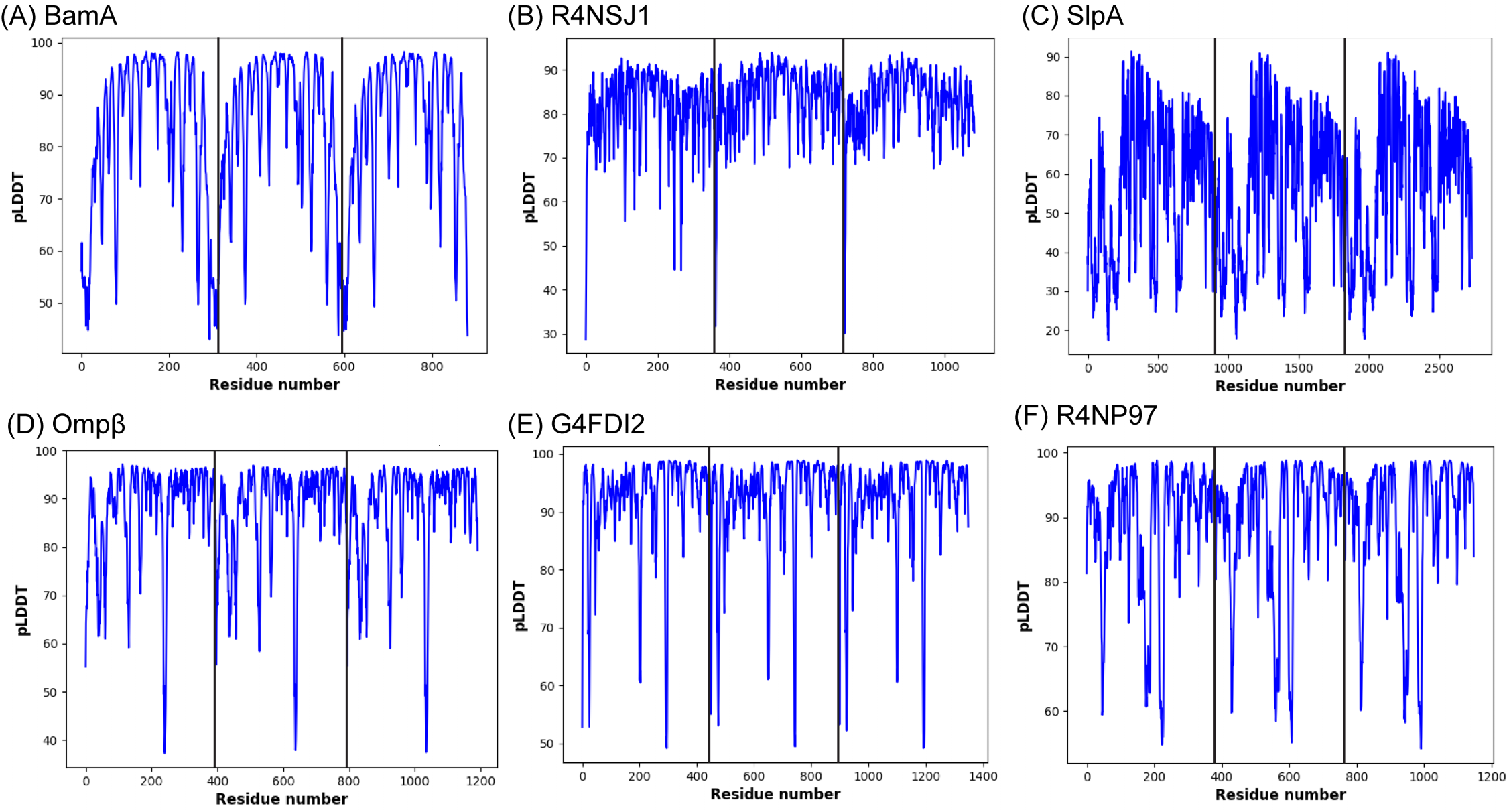
Per residue confidence plots (pLDDT) for structures predicted with AlphaFold-Multimer. The per residue confidence plot (pLDDT) for the model with the highest confidence depicted in Figure S3 is shown. Subunits for trimeric prediction are demarcated with solid black lines.

