## Supplemental Text for "The cell envelope of *Thermotogae* suggests a mechanism for outer membrane biogenesis"

**Extended Materials and Methods**

*Protein extraction and analysis*

For MS-based protein analysis, the separated IM and toga fractions were resuspended in 6x Laemmli sample buffer diluted in wash buffer and boiled for 10 min at 95 °C. Samples were run 5 mm into a 10% SDS-PAGE gel. The gel was stained with Coomassie and destained in water without exposure to heat. Gel bands were excised in small pieces of about 1 $mm^3$ using a scalpel blade. The gel bands were washed 3 times for 15 min in 50 mM ammonium bicarbonate/ acetonitrile (50:50, v/v), rinsed in 100% acetonitrile and incubated for 20 min in fresh 100% acetonitrile. After being air dried, proteins were reduced with dithiothreitol (DTT; 10 mM in 100 mM ammonium bicarbonate) for 30 min at 56 °C and alkylated with iodoacetamide (IAA; 50 mM in 100 mM ammonium bicarbonate) for 30 min in the dark at room temperature. The gel bands were then rehydrated with a trypsin solution (Promega; 0.02 µg/µL in 40mM ammonium bicarbonate and 10% acetonitrile) and incubated for 2 hrs on ice. The excess of trypsin solution was removed, and trypsin buffer (40 mM ammonium bicarbonate 10% acetonitrile) was added to cover the gel pieces. Trypsin digestion was performed for 16 h at 37 °C. After the digestion, the supernatant (tryptic peptides) was transferred into a new tube containing 5 µl of extraction solution (acetonitrile/water/10 % trifluoroacetic acid (60:30:10, v/v)). Tryptic peptides were extracted twice from the gel bands by incubation in the extraction solution and vortexing for 10 min. The extracted peptides were combined into the same Eppendorf tube. Samples were then lyophilized and resuspended in 10 µl of 1% formic acid in water. Tryptic peptides were analyzed on an Orbitrap Fusion Lumos Tribrid mass spectrometer (ThermoFisher Scientific) operated with Xcalibur (version 4.0.21.10) and coupled to a Thermo Scientific Easy-nLC (nanoflow Liquid Chromatography) 1200 system. Tryptic peptides were loaded onto a C18 trap (75 um x 2 cm; Acclaim PepMap 100, P/N 164946; ThermoFisher Scientific) at a flow rate of 2 µl/min of solvent A (0.1% formic acid in LC-MS grade water). Peptides were eluted using a 45 min gradient from 5 to 40% (5% to 28% in 40 min followed by an increase to 40% B in 5 min) of solvent B (0.1% formic acid in 80% LC-MS grade acetonitrile) at a flow rate of 0.3 µL/min and separated on a C18 analytical column (75 um x 50 cm; PepMap RSLC C18; P/N ES803; ThermoFisher Scientific). Peptides were then electrosprayed using 2.1 kV voltage into the ion transfer tube (300 °C) of the Orbitrap Lumos operating in positive mode. The Orbitrap first performed a full MS scan at a 120,000 full width half maximum (FWHM) resolution to detect the precursor ion having a *m/z* between 375 and 1,575 and a +2 to +7 charge. The Orbitrap AGC (Auto Gain Control) and the maximum injection time were set at 4 x $10^5$ and 50 ms, respectively. The Orbitrap was operated using the top speed mode with a 3 sec cycle time for precursor selection. The most intense precursor ions presenting a peptidic isotopic profile and having an intensity threshold of at least 5,000 were isolated using the quadrupole and fragmented with HCD (30% collision energy) in the ion routing multipole. The fragment ions ($MS^2$) were analyzed in the ion trap at a rapid scan rate. The AGC and the maximum injection time were set at 1 x $10^4$ and 35 ms, respectively, for the ion trap. Dynamic exclusion was enabled for 30 sec to avoid of the acquisition of same precursor ion having a similar m/z (plus or minus 10 ppm).

The Lumos raw data files were converted into Mascot Generic Format (MGF) using RawConverter (v1.1.0.18; The Scripps Research Institute) operating in a data dependent mode. Monoisotopic precursors having a charge state of +2 to +7 were selected for conversion. This MGF file was used to search against the *T. maritima* strain MSB8 proteome (Uniprot ID 243274) using Mascot algorithm (Matrix Sciences; version 2.7). Search parameters for MS data included trypsin as enzyme, a maximum number of missed cleavage of 1, a peptide charge equal to 2 or higher, cysteine carbamidomethylation as fixed modification, methionine oxidation as variable modification and a mass error tolerance of 10 ppm. A mass error tolerance of 0.6 Da was selected for the fragment ions. Only peptides identified with a score having a confidence higher than 95% were kept for further analysis. The Mascot data files were imported into Scaffold (v4.3.4, Proteome Software Inc) for comparison of different samples based on their mass spectral counting. Protein localization was predicted using pSORTb (1) and tertiary structures were predicted using RoseTTAFold (2).

*Lipid extraction and analysis*

Whole membrane, IM, and toga fractions were analyzed for their lipid composition using liquid chromatography-mass spectrometry (LC-MS). Briefly, 1.5 mL of a solvent composed of methanol:acetonitrile:water (2:2:1, vol:vol:vol) was added to each of the three glass tubes corresponding to the individual fractions. The mixture underwent sonication for 45 minutes to homogenize the samples. Technical replicates of each fraction were generated by splitting each fraction into three 400 µL aliquots into 2 mL Eppendorf vials. This was followed by subjecting the samples to three freeze/thaw (F/T) cycles for protein precipitation. One given cycle involved placing the samples in liquid $N_2$ for 1 minute prior to sonication in an ice-bath for 15 minutes. After completing the F/T cycles the samples were left overnight in a -20 °C freezer. Afterwards, 1.2 mL of MTBE was added to each sample, followed up with vortexing and repetition of the previous step, except using 0.4 mL $H_2O$. Subsequently, all samples underwent centrifugation at 14,000 rpm and 4 °C for 15 minutes. After centrifugation, the upper layer, containing the lipids, was transferred into new 1.5-mL Eppendorf vials. To dry the lipid samples, the solvent was evaporated using a vacuum concentrator at 4 °C overnight. A total of 150 µL of isopropanol:acetonitrile (1:1, vol:vol) was added to reconstitute the dried residue. The reconstituted solution was vortexed for 30 s and centrifuged at 14,000 rpm at 4 °C for 15 min. The resulting supernatants were transferred to glass inserts for LC-tandem MS (LC-MS/MS) analysis. Lastly, the QC sample was made by pooling equal aliquots (20 µL) of each sample together.

Lipid profiling of cellular membranes was carried out using an Agilent 1290 Infinity II ultra-high performance liquid chromatography (UHPLC) system (Agilent Technologies) coupled with Bruker Impact II electrospray-ionization quadrupole time-of-flight (ESI-QTOF) mass spectrometer (Bruker Daltonics). LC separation was performed using a reversed phase Acquity UPLC BEH $C_{18}$ column (1.0 × 100 mm, 1.7 µm, Waters), which was maintained at 25 °C. MS detection was performed in both ESI positive (ESI (+)) and ESI negative (ESI (−)) modes in two separate acquisitions. For ESI (+), mobile phase A was acetonitrile/water (6:4, v/v, pH = 4.8 adjusted by 0.1% formic acid) containing 5 mM ammonium acetate, and B was isopropanol/acetonitrile (9:1, v/v). For ESI (−), acetonitrile/water (6:4, v/v, pH = 9.8) and isopropanol/acetonitrile (9:1, v/v) were used as mobile phases A and B, respectively, with only mobile phase A containing 5 mM ammonium acetate. The gradient elution program for ESI (-) was as follows: 0 min, 5% B; 20 min, 95% B; 23 min, 95% B; 24 min, 5% B; 33 min, 5% B. For ESI (+) the gradient elution program was the same as ESI (-), with the exception of the last run at 35 min. The flow rate was set at 0.1 mL/min, and the injection volumes were optimized as 6 µL for ESI (-) and 3 µL for ESI (+). For the MS, capillary voltage was set at 4,500 V for ESI (+) and 3,600 V for ESI (−). The nebulizer gas was set at 1.0 bar, with drying gas flow rate of 6 L/min and source temperature of 220 °C. Data-dependent acquisition (DDA) mode was applied to collect both MS1 and MS2 spectra. A programed injection of 2 µL sodium formate (250 mM) at 25 min was used for internal mass calibration. HPLC-grade methyl tert-butyl ether (MTBE) was purchased from Merck. All the other high purity solvents and chemicals were purchased from ThermoFisher Scientific.

All the raw data were first calibrated using Bruker Compass Data Analysis (version 4.4) and then converted to ABF format with Abf Converter software. Lipid identification was completed on MS-DIAL 4.7 tool (3) based on mass accuracy, isotope ratio, retention time along with MS/MS similarity against publicly available libraries. MS-DIAL 4.7 can be downloaded at the PRIMe website (http://prime.psc.riken.jp/). Raw lipidomics data were collected in positive and negative ion mode, after intensity correction and normalization (post QC calibration). QC calibration was done to obtain more accurate fold changes. The relative standard deviation (rsd) of the QC injections (at the beginning, middle and end of the injection sequence) were below the threshold of 0.25. To measure enrichment of lipid species, fold changes (FC) of 0.67 and 1.5 were used as thresholds for the IM and toga, respectively. For example, FC were calculated as IM/toga ratio, thus FC of 0.67 indicated lipid enrichment in the toga, whereas FC of 1.5 indicated lipid enrichment in the IM fraction.
